# Expression dynamics of ARGONAUTE proteins during meiosis in Arabidopsis

**DOI:** 10.1101/2021.10.08.462716

**Authors:** Cecilia Oliver, German Martinez

**Affiliations:** Department of Plant Biology, Uppsala BioCenter, Swedish University of Agricultural Sciences and Linnean Center for Plant Biology, Uppsala, Sweden

## Abstract

Meiosis is a specialized cell division that is key for reproduction and genetic diversity in sexually reproducing plants. Recently, different RNA silencing pathways have been proposed to carry a specific activity during meiosis, but the pathways involved during this process remain unclear. Here, we explored the subcellular localization of different ARGONAUTE (AGO) proteins, the main effectors of RNA silencing, during male meiosis in *Arabidopsis thaliana* using immunolocalizations with commercially available antibodies. We detected the presence of AGO proteins associated with posttranscriptional gene silencing (AGO1, 2 and 5) in the cytoplasm or the nucleus, while AGOs associated with transcriptional gene silencing (AGO4 and 9) localized exclusively in the nucleus. These results indicate that the localization of different AGOs correlates with their predicted roles at the transcriptional and posttranscriptional levels and provide an overview of their timing and potential role during meiosis.

## Introduction

Meiosis is a special type of cell division where one round of DNA synthesis is followed by two rounds of cell division, segregating homologous chromosomes during the first division and sister chromatids at the second division (Marston et al. 2004, Mercier et al. 2015). This process is key for the production of gametes and the reshuffling of the genetic information during sexual reproduction (Bolcun-Filas et al. 2018). The mechanisms regulating meiosis have been widely studied at the cellular, genetic, and molecular levels in a variety of organisms. In plants, more than 90 genes have been identified comprising different meiotic processes that include double-strand break (DSB) formation, chromosome segregation or meiotic recombination (Huang et al. 2019a). Intriguingly, in the recent years it has been revealed that several of these processes involve the RNA silencing machinery (Oliver et al. 2016, Underwood et al. 2018, Wei et al. 2012). Different RNA silencing pathways are active during meiosis (Huang et al. 2020, Huang, et al. 2019a, Yelina et al. 2015). The miRNA affects chromatin condensation and the number of chiasmas, while the RNA-directed DNA methylation (RdDM) pathways affects chromatin condensation, the number of chiasmas and chromosome segregation (Oliver et al. 2017, Oliver, et al. 2016). Moreover, the RdDM pathway protects euchromatic regions from meiotic recombination (Yelina, et al. 2015). Additionally, *Arabidopsis* a non-canonical RNA silencing pathway plays a role in double-strand break repair (Wei, et al. 2012). Moreover, meiocyte-specific sRNAs between 23-24 nts are positively correlated with genes that have a meiocyte-preferential expression pattern (Huang, et al. 2019a), which could correlate with the observed role of DNA methylation in the regulation of gene expression in meiocytes (Walker et al. 2018). ARGONAUTE (AGO) proteins are the effectors of the different RNA silencing pathways and have dedicated members that act at the posttranscriptional or transcriptional levels. Here, we analyze the subcellular localization of the main AGO proteins in Arabidopsis during the different meiosis stages, which provides a confirmation of their activity during this process.

## Materials and Methods

### Plant material

Plants used for immunolocalization analysis were grown in a phytotron under long day conditions (16-hour light/8-hour dark photoperiod), at 24-25 °C and 45% relative humidity.

### Bioinformatic analysis

sRNA data was downloaded from the SRA repository project number PRJNA510650 (Huang et al. 2019b). sRNA alignments were performed using bowtie (Langmead et al. 2009) with the following parameters –t –v2 that allows 2 mismatches to the alignments. Alignment files were subsequently analyzed in Galaxy (Afgan et al. 2018). For sRNA categorization as miRNAs, sRNA libraries were aligned to individual indexes generated for each genomic category and compared total sRNAs mapping to the TAIR10 chromosome sequences. The miRbase version 22.1 (https://www.mirbase.org/) was used for miRNA alignments (Kozomara et al. 2019). Transcriptomic data corresponds to the CATMA arrays data from GEO accessions GSE10229 and GSE13000 (Libeau et al. 2011). CATMA array data was extracted using the CATdb database (http://urgv.evry.inra.fr/cgi-bin/projects/CATdb/catdb_index.pl) were normalized data was extracted for both GSE10229 (http://urgv.evry.inra.fr/cgi-bin/projects/CATdb/consult_expce.pl?experiment_id=195) and GSE13000 (http://urgv.evry.inra.fr/cgi-bin/projects/CATdb/consult_expce.pl?experiment_id=46).

### Cytology

Immunolocalization on meiotic nuclei were carried out by squash technique as was previously described by Manzanero et al. (2000) with some modifications (Oliver et al., 2013). Two bioreplicates constituted by young flower buds from five different plants, were analyzed. Young flower buds were fixed for 20 min in freshly prepared 4 % (w/v) paraformaldehyde, 0.1 % (v/v) Triton X-100 in phosphate-buffered saline (PBS, pH 7.3). Flower buds were then washed at room temperature for 30 min in PBS that was changed twice. Buffer was removed before incubation at 37ºC during 20–40 min with an enzyme mixture of 1 % pectinase, 1 % cellulase and 1 % cytohelicase (w/v) (Sigma), dissolved in PBS. Buds, immersed in a small volume of PBS, were transferred to slides with a Pasteur pipette, macerated with a needle and squashed between a glass slide and cover slip. After freezing in liquid nitrogen, the cover slips were removed and the slides were transferred immediately into PBS. Prior to immunostaining experiments the slides were washed twice in PBS, 0.1 % (v/v) Triton X-100 for 5 min each. To avoid non-specific antibody binding, slides were incubated for 30 min in PBS with 1 % BSA (w/v) and 0.1 % Triton X-100 at room temperature. The incubation with the primary antibody was carried out in a humidified chamber. The primary antibodies used were rabbit anti-AGO1 (1:200 AS09 527), -AGO2 (1:100, AS13 2682), -AGO5 (1:100, AS10 671), -AGO4 (1:100, AS09 617), -AGO6 (1:50, AS10 672), -AGO9 (1:100, AS10 673) and -AGO10 (1:50, AS15 3071) antibodies from Agrisera. All the primary antibodies were diluted in PBS, 1 % BSA, 0.1 % Triton X-100. After overnight incubation at 4ºC and washing for 15 min in PBS with 0.1 % Triton X-100, the slides were incubated for 1 h at room temperature with goat anti-rabbit IgG H&L Alexa Fluor 568 conjugated (1:200; ab175471; Abcam) diluted in 1 % BSA, 0.1 % Triton X-100 in PBS. Slides were then washed in PBS, 0.1 % Triton X-100, before they were stained the DAPI, 1 μg/ml during 20-30 min and finally mounting with antifading medium (0.2% n-propyl Gallete, 0.1% DMSO, 90% glycerol in PBS). Fluorescent signals were observed using an epifluorescence microscope Zeiss AxioScope A1. Images were captured with AxioCam ICc5 camera and were analyzed and processed with ImageJ and Affinity Photo software.

## Results

To discern the level of expression of RNA silencing components in meiocytes, we analyzed their expression from publicly available microarray datasets (Libeau, et al. 2011) (Figure 1 and Supplementary Methods). Overall, several components from the RNA silencing pathways were preferentially expressed in meiocytes compared to somatic tissues (Figure 1A), including the AGO proteins AGO4, 5 and 10, the Dicer-like (DCL) proteins DCL1, 3 and 4 or the sRNA methyltransferase HEN1. This indicated that different PTGS (AGO5, DCL1 and DCL4) and TGS (AGO4 and DCL3) pathways might be especially active during meiosis. Previous analysis (Huang, et al. 2019a) have shown that TE-derived sRNAs accumulate to relatively high levels in meiocytes and that certain miRNAs like miR845 are active before the microspore stage (Borges et al. 2018). Although miRNAs were not globally enriched in meiocytes (Figure 1B), several miRNAs were strongly upregulated including miR839, miR780.2, miR780.1, miR157, miR172, miR166 and miR860, which are important regulators of several transcription factor families (Figure 1C, Supplementary Figure 1 and Supplementary Table 2). In summary, transcriptomic and sRNA sequencing analysis supported the notion that the RNA silencing machinery might have a meiocyte-specific activity.

**Figure 1.**
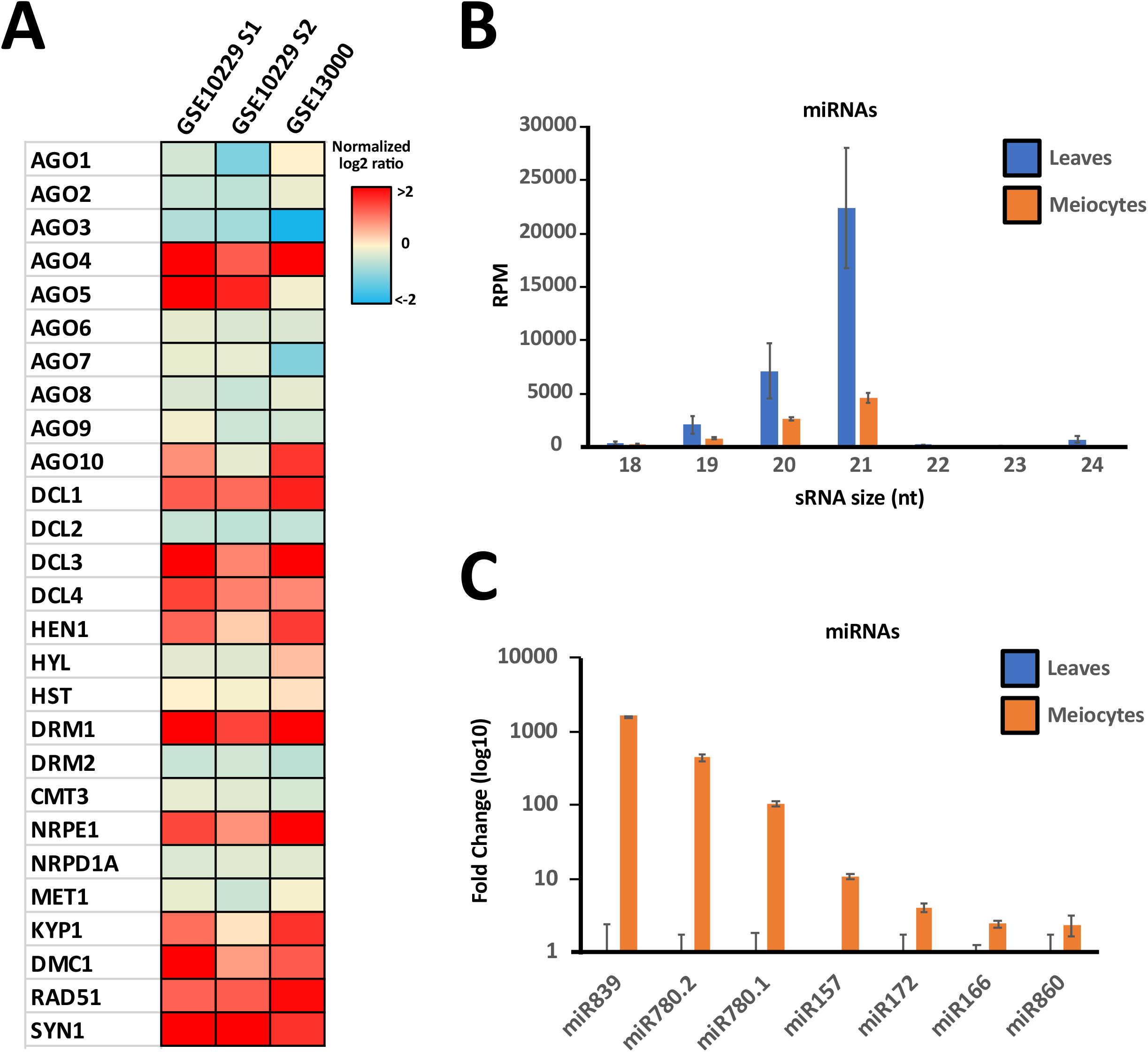
Analysis of the expression in meiocytes of different RNA silencing and epigenetic pathways components and analysis of miRNA accumulation in meiocytes. **A**. Heat map of the expression values of RNA silencing and epigenetic pathways components in meiocyte microarray experiments. Expression values are represented as the normalized log2 ratio of the comparison meiocyte/control tissue. **B**. Global accumulation of miRNAs in leaves and meiocytes samples from public datasets normalized to reads per million. **C**. Accumulation values of miRNAs enriched in meiocyte sRNA libraries. Enrichment was considered only for miRNAs accumulating more than 2-fold in meiocytes and with a p-value<0.05.

Although transcriptomic analysis is important to infer the activity of the different RNA silencing pathways in meiocytes, this analysis provides a steady image of this tissue and ignores, for example, its dynamism during meiosis. To understand the subcellular localization and dynamics of the different AGO proteins during meiosis, we performed immunolocalizations of the AGO proteins that had commercially available antibodies (Agrisera, AGO1, 2, 4, 5, 6, 9 and 10, Figure 2 and Supplementary Methods). During meiosis all AGOs but AGO6 and AGO10 could be detected. In detail, AGO1 and its paralogs AGO2 and AGO5 displayed a similar localization and expression pattern during the first meiotic stages (Figure 2A, 2B, 2C). The three proteins were located mainly in the cytoplasm, similar to their localization in somatic tissues (Bologna et al. 2018, Ye et al. 2012). From the leptotene to the diplotene stage these three AGO proteins formed cytoplasmic granules (Figure. 2A1, 2B1, 2C1). In somatic tissues, cytoplasmic bodies are involved in the degradation and translation arrest of mRNAs (Maldonado-Bonilla 2014). In mammals, AGO proteins localize in P-bodies where they mediate the translational repression of their target mRNAs (Liu et al. 2005). The localization pattern observed for AGO1, 2 and 5 might indicate a similar role of RNA silencing in the posttranscriptional regulation of mRNAs, a process that is known to take place in other organisms like mammals (Yao et al. 2015).

**Figure 2.**
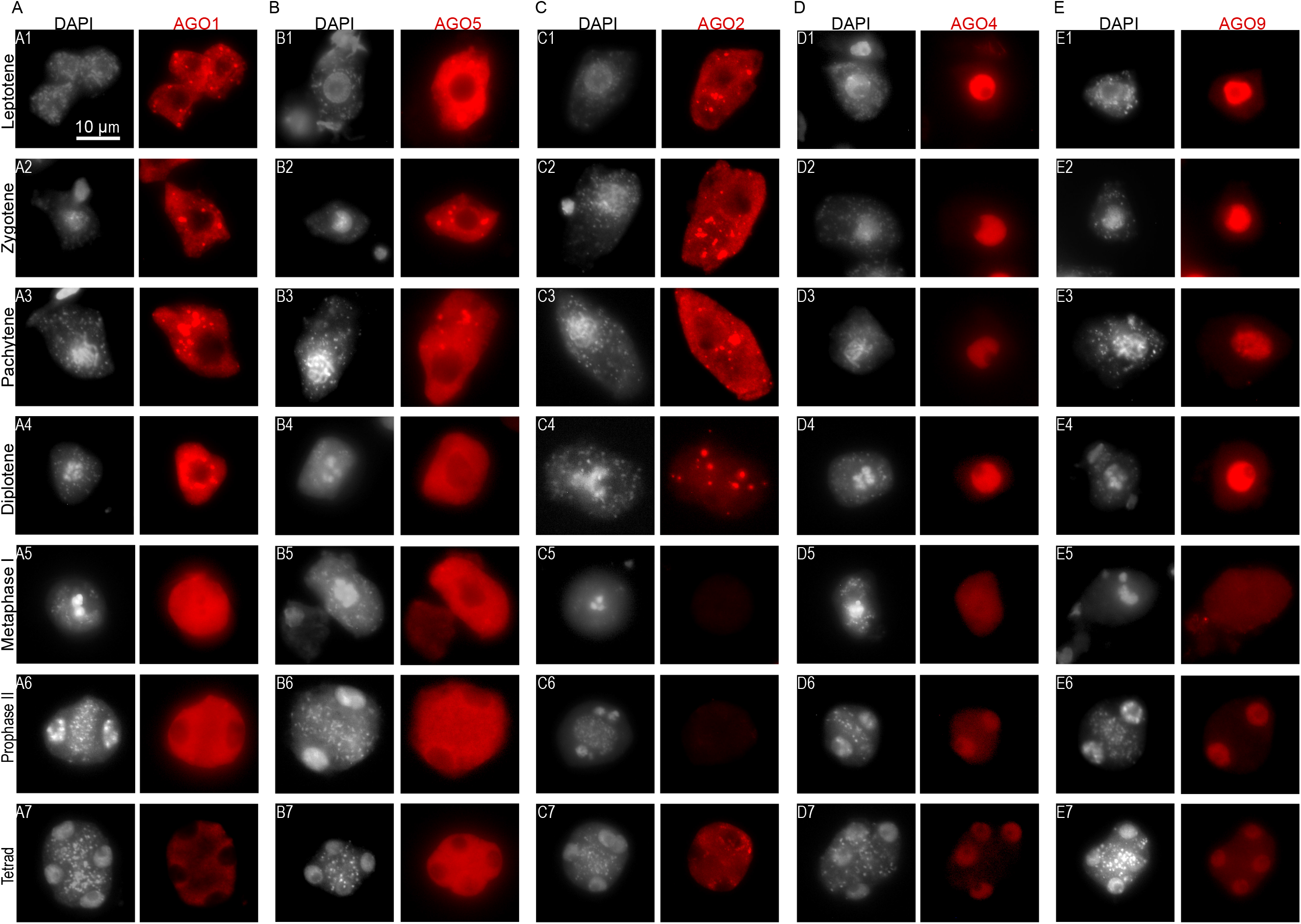
Immunolocalization of AGO1 (A), AGO5 (B), AGO2 (C), AGO4 (D) and AGO9 (E) at different representative meiotic stages in Arabidopsis meiocytes. Leptotene (A1, B1, C1, D1, E1); Zygotene (A2, B2, C2, D2, E2); Pachytene (A3, B3, C3, D3, E3); Diplotene (A4, B4, C4, D4, E4); Diakinesis (B5), Metaphase I (A5, C5, D5, E5) Prophase II (A6, B6, D6, E6); Metaphase II (C6); Tetrad (A7, B7, C7, D7, E7). Immunostaining with antibodies is shown in red, counterstaining with DAPI is shown in grey. Bar indicates 10 µm.

Despite the similarities between the accumulation during meiosis, AGO1, 2 and 5, they showed differences in their dynamics during meiosis. For example, AGO1 condensates around the nuclear envelope at diplotene (Figure 2A4) but after this stage, it showed a disperse accumulation (Figure 2A5). This location during cell division could be related with the known AGO1 association with the endoplasmic reticulum (Li et al. 2013), as when the nuclear envelope disassembles it reorganizes in vacuoles around the bivalents (Marston, et al. 2004, Mercier, et al. 2015). AGO5 displayed a similar pattern of subcellular localization to AGO1, although its localization at cytoplasmic bodies disappeared at diplotene (Figure 2B4). On the other hand, AGO2 showed a dual localization in the cytoplasm and in the nucleus (Figure 2C1-4) and was not detectable after metaphase I (Figure 2C5-6). Both its nucleocytoplasmic localization and timing of expression are in line with its known role in double strand break (DSB) repair, which takes place during the first meiotic stages (Oliver et al. 2014, Wei, et al. 2012). Nevertheless, AGO2 expression pattern was recapitulated after the second meiotic division (Figure 2C7), indicating that it might serve other roles in parallel to its function in DSB repair during meiosis.

On the other hand, the TGS/RdDM-associated AGO proteins, AGO4 and AGO9, were located in the nuclei during all meiotic stages (Figure 2D and E). Exceptionally, at metaphase I, when the nuclear envelope dissolves, both proteins showed a dispersed accumulation. This is in accordance with the known role of the RdDM pathway in regulating DNA methylation during meiosis (Walker, et al. 2018). Meiocytes have the lowest CHH methylation values of all the reproductive nuclei analyzed, but its activity is needed for the regulation of gene expression (Walker, et al. 2018). We detected a low accumulation of AGO4 and 9 after metaphase I (Figure 2D5-6 and 2E5-6), which might partially cause this reduction in CHH methylation. Interestingly, we observed that AGO9 displayed a localization pattern compatible with a preference for heterochromatic regions at pachytene. This localization might explain the known role of AGO9 on the dissolution of interlocks during meiosis (Oliver, et al. 2014).

## Discussion

In summary, our results provide an overview of the subcellular localization, timing and potential role of different RNA silencing pathways during meiosis. Furthermore, our work complements previous analysis that analyzed RNA silencing activity in meiocytes, and opens the door for future molecular analysis of the specific role of AGO proteins during specific meiosis stages, which are technically challenging at the moment.

## Author contribution statement

C.O and G.M. design the experiments and wrote the manuscript. C.O. performed the experiments and analyzed the data. G.M. analyzed the bioinformatic data.

## Acknowledgements

The authors thank SLU, the Carl Tryggers Foundation (CTS 17-305 and CTS 18-251), the Swedish Research Council (VR 2016-05410) and the Knut and Alice Wallenberg Foundation (KAW 2019-0062) for supporting research in the Martinez group. The data handling was enabled by resources provided by the Swedish National Infrastructure for Computing (SNIC) at UPPMAX partially funded by the Swedish Research Council through grant agreement no. 2018-05973.

## Figure legends

**Supplementary Figure 1**. Predicted and confirmed targets of miRNA families significantly upregulated in meiocytes.

**Supplementary Table 1**. Raw values of normalized log2-ratio expression values for selected genes in meiocytes microarray data.

**Supplementary Table 2**. Raw values of miRNA accumulation in meiocytes and leaf sRNA libraries.

