## Supplementary Figure 1 for "Expression dynamics of ARGONAUTE proteins during meiosis in Arabidopsis"

**miR839**

```
 miRNA     19 CUUGCUACUUU-CCAACCAU 1
              :::.::::::: ::::::::
Target   1330 GAAUGAUGAAAUGGUUGGUA 1349
```

**AT2G33710**

ERF (ethylene response factor) subfamily B-4 of ERF/AP2 transcription factor family.

**Predicted**

**miR780**

```
 miRNA     21 UACGGUCUAUAAGUGCUUCUU 1
              :::.: ::::::::.:::::
Target   1874 AUGUCUGAUAUUCAUGAAGAU 1894
```

**AT5G41610**

member of Putative Na+/H+ antiporter family

**Confirmed**

**miR157**

```
 miRNA     21 CACGAGAGAUAGAAGACAGUU 1
              ::::::::: :::::::::::
Target   2368 GUGCUCUCUCUCUUCUGUCAA 2388
```

**AT1G27370, AT5G43270, AT1G27360, AT2G4220, AT3G57920, AT1G69170...**

Squamosa promoter binding protein-like

**Confirmed**

**miR172**

```
 miRNA     21 UACGUCGUAGUAGUUCUAAGA 1
              ::::::::::::::.::::::
Target   1647 AUGCAGCAUCAUCAGGAUUCU 1667
```

**AT5G60120, AT4G36920, AT5G67180, AT2G28550 ...**

AP2 family transcription factor

**Confirmed**

**miR166**

```
 miRNA     21 GGAGCUCGGU-CUGUUGUCAGG 1
              :.:.::::.: ::..:..::::
Target    354 CUUUGAGCUAUGAUGAUGGUCC 375
```

**AT5G60690, AT2G34710, AT2G46685...**

HD-ZIPIII family members

**Confirmed**

**miR860**

```
 miRNA     21 UAUGUAUCAGGUUAGAUAACU 1
              :::.::: ::::::.:.::::
Target    145 AUAUAUAUUCCAAUUUGUUGA 165
```

**AT3G12640, AT1G24967, AT3G33139**

RNA-binding proteins, transposable elements

**Predicted/ Confirmed**
